# Simulating serial-target antibacterial drug synergies using flux balance analysis

**DOI:** 10.1101/030791

**Authors:** Andrew S. Krueger, Christian Munck, Gautam Dantas, George M. Church, James Galagan, Joseph Lehár, Morten O.A. Sommer

**Author notes:** Equal contribution. Corresponding authors JL, MOAS).

## Abstract

Flux balance analysis (FBA) is an increasingly useful approach for modeling the behavior of metabolic systems. However, standard FBA modeling of genetic knockouts can not predict drug combination synergies observed between serial metabolic targets, even though such synergies give rise to some of the most widely used antibiotic treatments. Here we extend FBA modeling to simulate responses to chemical inhibitors at varying concentrations, by diverting enzymatic flux to a waste reaction. This flux diversion yields very similar qualitative predictions to prior methods for single target activity. However, we find very different predictions for combinations, where flux diversion, which mimics the kinetics of competitive metabolic inhibitors, can explain serial target synergies between metabolic enzyme inhibitors that we confirmed in *Escherichia coli* cultures. FBA flux diversion opens the possibility for more accurate genome-scale predictions of drug synergies, which can be used to suggest treatments for infections and other diseases.

## Introduction

Microbial infections are a major burden on societies, and the emergence of drug-resistant bacteria poses an increasing threat to human welfare. Drug combinations can overcome resistance by creating new therapeutic avenues inaccessible to single target drugs [1-3] or eliminating functional redundancies exhibited by robust biological networks [4]. However, although combinations are increasingly the standard of care for many bacterial infections [5,6] the complexity of microbial biology and the vast number of possible target combinations makes finding new effective drug combinations challenging.

Systems biology may provide a solution to this challenge [7-9], by modeling microbial systems as complex networks of interacting components. Dynamic models of microbial function under drug treatment [10,11] can provide detailed and accurate representations of phenotypes, but the scale of such models is limited by the scarcity of kinetic molecular reaction rate measurements. Graph-theoretic networks of metabolic interactions found in KEGG or Metacyc [4,12,13] can address the true scale of microbial biology, but are limited to static representations with only limited relevance to drug response phenotypes.

A successful approach towards genome-scale modeling is Flux Balance Analysis (FBA) [14-16], which utilizes reaction stoichiometry to model metabolic capabilities at steady state (Fig 1A). By integrating the properties of metabolic networks into a single growth phenotype, FBA enables predictions of enzymatic gene essentiality and even genetic interactions. FBA approaches have been used to predict the nutrient dependent metabolic phenotype of gene knockouts (Fig 1B) [15,17], and have been successfully applied to model the combined effects of double knockouts in microbial systems [18]. Recently, integration of proteome structure has allowed for the prediction of the temperature dependence of metabolic reactions on a genome wide scale [19]. Several methods have also been developed to incorporate gene expression and other high throughput data to constrain fluxes through particular reactions predicting metabolic states corresponding to specific gene expression states; E-flux in particular has been used to predict the impact of drugs given expression data [20-24]. Finally, incorporation of gene expression networks and protein translation processes has enabled a mechanistically detailed description of cellular trade-offs occurring during various growth phases and nutrient limitations[25-29]. However, none of the above methodologies models the effects of drug dosing. The continuous responses to varying inhibitor concentrations are especially critical for identifying and interpreting drug combination effects[10].

**Fig 1.**
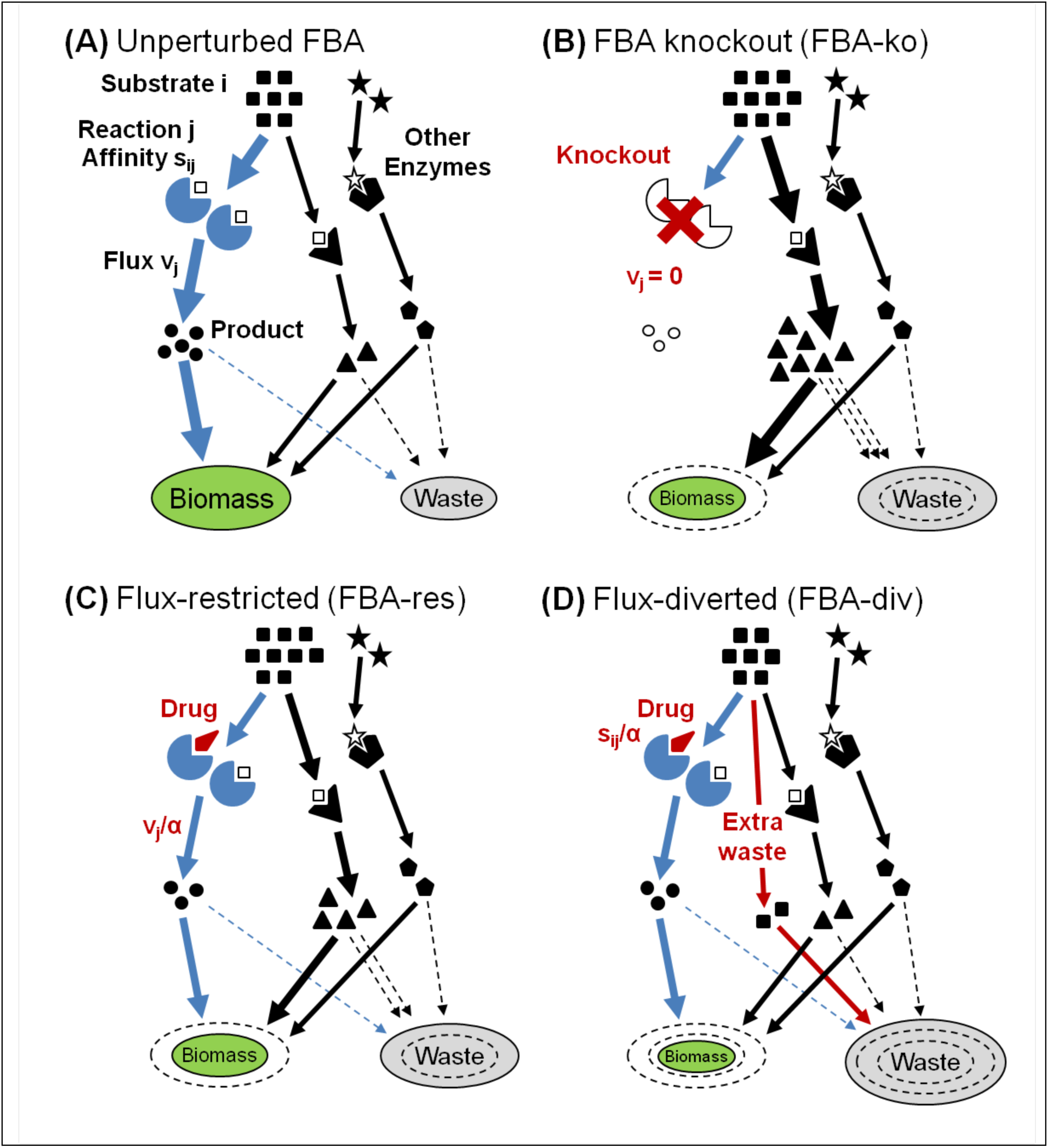
Simulations of inhibited FBA metabolism. (**A**) Cartoon of a target enzyme “j” which acts on substrate “i” at a steady-state velocity **v**_j_. Other enzymes may compete for the same substrate, and downstream reactions will convert all products to “biomass” flux or unproductive “waste” that is degraded or exported. (**B**) When the target is knocked out by setting **v**_j_=0, substrate backlog increases flow through other reactions, and increased waste rates reduce biomass. (**C**) FBA-res reduces the target velocity by a scalar factor α, causing a partial knockout effect. (**D**) For FBA-div, the reaction’s affinity is reduced by scaling its stoichiometric coefficient **s**_ij_, diverting the backlog to waste. This yields stronger effects than FBA-res, reducing effects on the rest of the network.

Here we have extended FBA modeling to simulate drug effects over multiple doses. We explore new methods for simulating drug treatments with FBA, considering both direct target flux restriction[3] and a model that diverts flux to non-productive waste pathways (Fig 1). In flux restriction “FBA-res”, the flux through the target reaction is restricted by a variable scalar factor (Fig 1C), while in flux-diversion “FBA-div”, we instead divert flux to a waste reservoir (Fig 1D). The two methods yield similar results to knockout simulations for single agent effects, but have very different predictions for combinations. Only FBA-div predicts potent antibiotic synergies targeting metabolism.

## Materials and methods

### Simulations

The Escherichia Coli iAF1260 model created by Bernhard Palsson’s group at UCSD serves as our bacterial model in this work[30], which contains species-specific metabolic reactions, linked together in a network by substrates and products. For our simulations, we will assume bacterial growth on rich media with ample supplies of oxygen, glucose, ammonia, potassium, sulphur, and all amino acids.

All drug effect simulations were performed with an R-script using the R package Sybil (Systems Biology Library for R)[31] and the genome-scale reconstruction of Escherichia Coli. The “Ec_iAF1260_flux2” model was downloaded in xml format and added to the R workspace. After downloading, the growth flux is calculated using the built in optimization module “optimizeProb” with the algorithm set to “fba”. The theoretical framework for FBA-res models drug perturbations as a scalar restriction of flux through a targeted reaction. We implement FBA-res by reducing the flux bounds by a scalar alpha for each dose of every target. This creates a new drug perturbed model. The drug perturbed model is sent to optimizeProb with the algorithm set to “fba”. Growth inhibition is calculated with the unperturbed and perturbed growth fluxes. After each dose for combinations or single agents, we reset to the original model to implement the next perturbation. The theoretical framework for FBA-div models drug perturbations as a scalar diversion of flux, which depends on the magnitude of the perturbation (dose). We implement FBA-div by first adding waste reactions and waste metabolites to the model. Initially the waste metabolites do not belong to any reaction, and the waste reactions consume waste metabolites irreversibly. To simulate drug effects, the amount of metabolites produced by a targeted reaction are reduced by alpha and the remainder of mass is converted into a waste metabolite, which is now connected to the targeted reaction. Waste reactions consume any diverted waste metabolites. In the case of reversible reactions, two irreversible reactions are created with different waste metabolites, and alpha is applied to both reactions. This is the drug perturbed model. The drug perturbed model is sent to the optimization module “optimizeProb” with the algorithm set to “fba”. Growth inhibition is calculated with the unperturbed and perturbed growth fluxes. After each dose for combinations or single agents, we reset to the original model to implement the next perturbation.

Drug effects on a targeted metabolic reaction were implemented in the genome-scale model either by limiting the target flux (FBA-res) or limiting the amount of mass converted from substrate to product by diverting metabolic mass to a waste reaction (FBA-div). Inhibition values Inhib = 1 - *f*_treat_/*f*_wt_, where *f*_wt_ and *f*_treat_ are the simulated biomass flux rates for untreated and drug treated bacteria. In both cases the MOMA quadratic programming algorithm for simulating perturbed biological networks was implemented to find the change in biomass flux. The IC50 value for a reaction describes the amount by which reaction flux must be reduced to inhibit growth by 50%. For FBA-res, flux is directly restricted by a scalar drug concentration, and so the IC50 is the scalar that achieves a 50% growth inhibiting flux restriction. For FBA-div, flux is diverted to a waste reaction, and so the IC50 is the value that achieves 50% growth inhibiting flux diversion. First, a central perturbation of α_cent_ (where α_cent_=1+[Inhibitor]/K_i_ and for simplicity K_i_=1) was found for each target enzyme using a bisection search to yield half the inhibition level of a full FBA gene deletion (for enzymes showing no deletion phenotype, we used α=5,000,000 since this value reproduced synthetic lethal interactions in[12]. Simulated response curves were then generated for each enzyme, with five concentrations centered on this α_cent_ using 3x dilution steps of inhibitor concentration (α-1), covering a ~100x dynamic range (Fig 2A). For each pair of agents, we then generated dose matrices by simulating combined inhibitors at all pairs of those concentrations (Fig 2B).

**Fig 2.**
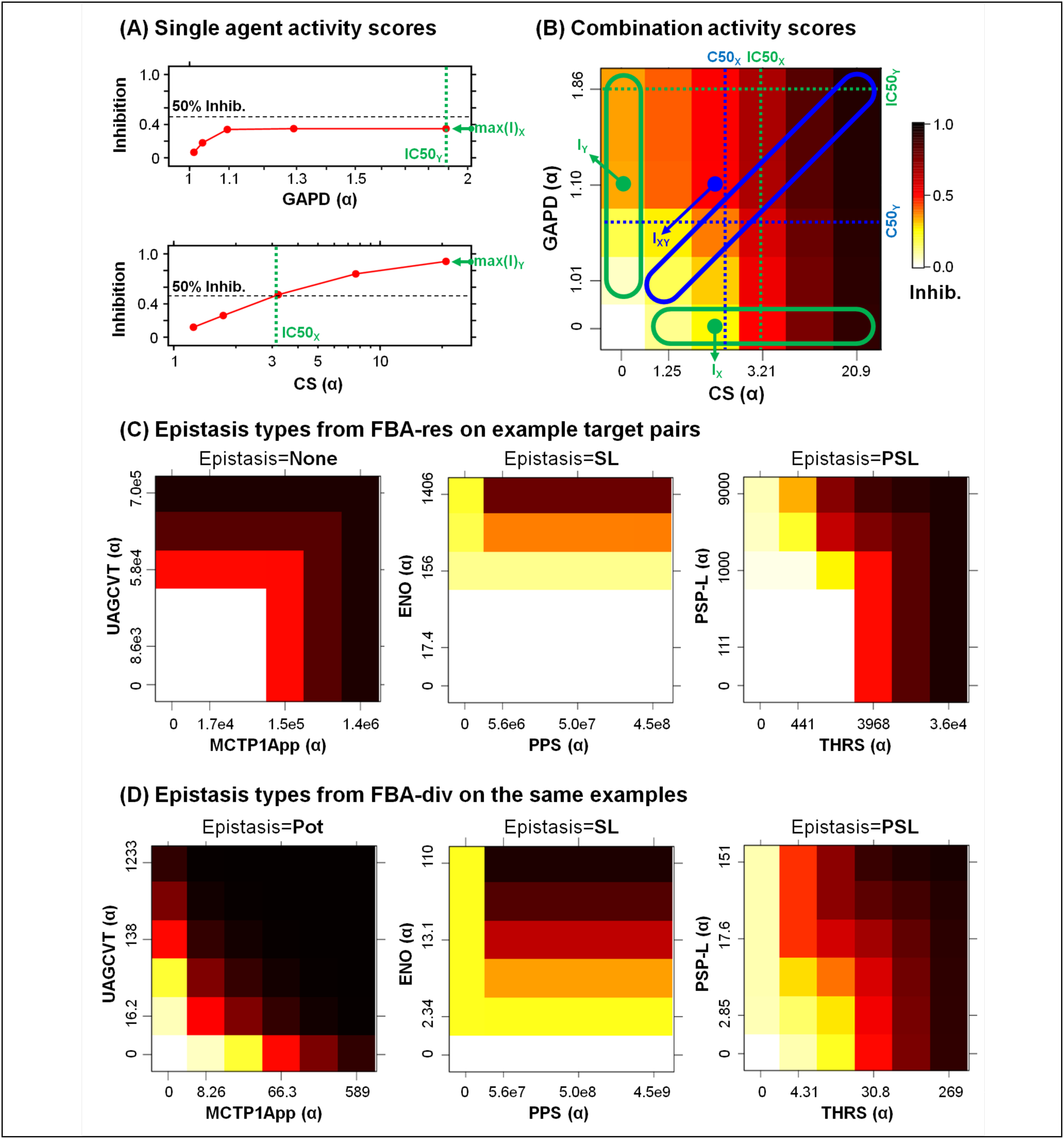
Activity and synergy measures for each combination. (**A**) Inhibition of each enzyme was simulated at five α concentrations, using 3-fold dilution intervals centered on the concentration that yielded half of the maximum simulated effect. Note that the 3x dilution steps were constant in simulated concentration α-1, so they are not always equal on these curve plots based on α. The maximum inhibition **max(I)** and 50% crossing concentration **IC50** was measured for each target (IC50 = top concentration if max(I)<0.5), and each combination was simulated at all 25 concentration pairs. (**B**) For combinations, we scored the maximum effect **max(I)** and synergy over Gaddum’s “best single agent” reference **max(ΔI)** = max{I_XY_ - max(I_X_,I_Y_)} across all inhibitions I_XY_. We also calculated “potency shift index” from the 50% inhibition crossing concentrations (C50_X_,C50_Y_) along the matrix diagonal (blue), where **SI50** = max(C50_X_/IC50_X_,C50_Y_/IC50_Y_). Epistasis types for (**C**) FBA-res and (**D**) FBA-div simulations on three example target pairs. Interactions were classified based on the single agent and combination effect and synergy, as either non-interacting “None”, synthetic lethality “SL” between inactive inhibitors, partial SL “PSL” involving one active agent, or potentiation “Pot” between two active inhibitors.

Complete analysis methods are provided as an R-project package, which includes the metabolic network matrix, the analysis code, and all supporting data files. Complete simulation results are also provided in S1 Table.

### Single agent effect, combination synergy score, and epistasis

We used simple metrics of single agent and combined inhibitor responses (Fig 2A), based on maximum inhibition levels max(I) or on 50% inhibitory concentrations IC50. Comparing max(I) values permits comparisons with standard FBA knockouts, while IC50s allow fuller use of the dose-responsiveness of inhibitors. The maximum inhibition max(I) and 50% crossing concentration IC50 was measured for each target, and if max(I)<0.5, IC50 was set to the top concentration.

Synergy was then determined for each dose matrix using either an effect difference max(ΔI) or “shift index” SI50, compared to the single agents (Fig 2B). The combination data I_XY_ (blue point) are compared to the corresponding single agent response values I_X_,I_Y_ (green points), and max(ΔI) = max{I_XY_ - max(I_X_,I_Y_) }, shows the Gaddum “best single agent” expectation[32]. All combination points were considered, with the largest difference across the matrix recorded as the score. Positive max(ΔI) values correspond to synergy (more effect than the better single agent at comparable doses). For dose shifting, the matrix diagonal (blue area) is compared to the two single agent curves (green areas). The combination’s 50% inhibitory crossing point along the diagonal has single agent component concentrations C50_X_ and C50_Y_, which are compared to the single agent 50% inhibitory concentrations IC50_X_ and IC50_Y_, to calculate SI50 = max(C50_X_/IC50_X_, C50_Y_/IC50_Y_). This SI50 indicates whether the combination shows synergistic (SI50<1) or antagonistic (SI50>1) dose shifting, relative to the Gaddum response surface. We also calculated a standard Loewe combination index CI50 = C_X_/IC50_X_ + C_Y_/IC50_Y_, which measures synergy relative to Loewe dose additivity[33], which is used to determine whether a combination outperforms a “drug-with-itself” sham combination (CI50=1 for dose-additive). All scores for simulated and experimental combinations are listed in S1 Table.

Simulated epistasis was classified (Fig 2C and D) based on the max(I) values for a combination and its single agents. If a combination’s max(I) < 0.05 or max(ΔI) < 0.05, we set epistasis to non-interaction “None”, corresponding to a Gaddum “best single agent” interaction model. Epistasis was then set to potentiation “Pot” if single agent max(I) >=0.5 for both agents, synthetic lethal “SL” if max(I) < 0.5 for both agents, or partial synthetic lethal “PSL” otherwise. To account for higher noise levels in experimental combinations, we used the same classification scheme, except that epistasis=None was called when max(I) < 0.25 or max(ΔI) < 0.25 for a combination. Epistasis calls are recorded in S1 Table.

### Proliferation assay

To experimentally validate the predicted drug interactions, 28 drug-pairs were tested against *E. coli* MG1655 (S3 Table). In order to explore the full interaction surface the drug pairs were tested in an 8×8 well 2d drug-gradient matrix in a 96 well plate, in addition to the drug matrix the plate also contained gradients for each of the sing drugs as well as negative and positive controls. For both the drug matrix and the single drug gradients the highest drug concentrations were 4 times above the minimal inhibitory concentration (MIC). The plates were inoculated with a 96 pin replicator and incubated 18-20 hours at 37° C. After incubation, OD600 was read using a BioTek H1 plate reader. Each plate was produced in five replicates. Raw data are presented in S4 Table, and response scores are in S1 Table.

## Results

### Flux balance analysis frameworks for modeling drug inhibition

Extensive networks of microbial metabolism have been constructed that link together thousands of species-specific metabolic reactions mined from the literature and online databases [34]. Simulating the behavior of microbial metabolism requires not only the stoichiometry captured by such networks, but also detailed kinetic parameters which are unknown for most reactions – even for well-studied microbes like *E. coli.* FBA addresses this limitation by replacing those parameters with linear fluxes through all the metabolic reactions, and using linear programming to derive steady-state flux values optimized on an objective function constructed from experimental abundances of nucleotides, amino acids and of anabolic metabolites [15,16,35]. FBA models optimize steady-state production of these essential building blocks, enforcing consistency with limits imposed by the network’s connectivity, flux limits and the conservation of mass between reactions.

FBA models have been used to successfully predict the effects of genetic knockouts [15,17]. In FBA, under steady state, a system of j=1…N reactions between i=1…M metabolites should satisfy Σ_j_ **s**_ij_ **v**_j_ = 0, where **v**_j_ are the reaction velocities and **s**_ij_ are stoichiometric coefficients that account for reaction affinities and connectivity. FBA models solve for the **v**_j_ that maximize simulated flux through the objective function, constrained by this mass conservation requirement and any v_j,min_<**v**_j_<v_j,max_ limits. To model gene essentiality using the standard knockout “FBA-ko” approach, the target enzyme’s reaction rate is set to **v**_j_=0, after which the model is re-optimized with a linear programming algorithm to maximize the objective function (Fig 1B and S1 Fig). Growth is represented by a biomass reaction which integrates the outputs from many metabolic pathways. A refinement to FBA applies “minimization of metabolic adjustment” (MOMA)[36], which requires the re-optimized reaction coefficients to minimize their distance from the unperturbed values, rather than seeking the maximum objective function flux consistent with the applied constraints. MOMA has been applied to *E. coli* and yeast metabolic networks and can successfully predict phenotypic responses to single[36] and double knockout experiments[18].

Extending FBA to drug perturbations, we consider two approaches: our previous “FBA-res” directly restricts the target flux while our new “FBA-div” diverts flux to non-productive waste. FBA-res reduces the velocity **v**_j_ through a targeted reaction by a scalar factor α (Fig 1C and S1 Fig). Thus, instead of FBA’s **v**_j_=0 constraint, we set **v**_j_➔**v**_j_/α and re-optimize the reaction fluxes. FBA-div modeling scales down the target’s stoichiometric affinity, rather than its maximum reaction rate (Figure 1D and S1 Fig). Specifically, FBA-div inhibits an enzyme by setting **s**_ij_➔**s**_ij_/α, for metabolites targeted by the inhibitor, before re-optimizing to the objective function. To conserve mass, we introduce transport reaction, **s**_i(N+1)_, with compensating stoichiometric coefficients **s**_i(N+1)_=1-Σ_j_ **s**_ij_/α that divert excess substrate to an infinite waste sink. We use standard FBA linear programming to solve the optimization problem in conjunction with our FBA-div methodology. In contrast to FBA-res, this waste diversion prevents other enzymes from increasing their reaction rates in response to target flux restriction. This usually predicts greater biomass reductions than FBA-res for the same level of target flux inhibition.

### Simulating drug epistatic interactions

To evaluate these approaches, we simulated combination effects using standard FBA-ko, FBA-res and FBA-div, applied to the iAF1260 model of *Escherichia coli* metabolism [30]. To explore mechanistic patterns, we chose 50 enzymes to cover synthetic lethal synergies and antagonisms found using FBA-ko [12] and sample key pathways in bacterial metabolism (S2 Table). Inhibition of each target was simulated at 5 “drug” concentrations using FBA-res and FBA-div, in each case estimating inhibition by the flux through the growth reaction in perturbed and unperturbed states (Methods and S1 Fig). We used the same methodology to generate combination response matrices across all 25 pairings of single drug concentrations. To score the simulation results, we used metrics of single agent and combined inhibitor responses, based on maximum inhibition levels or on 50% inhibitory concentrations IC50 (Fig 2A and B), and focused our analysis on the maximum effect max(I) and synergy score max(ΔI). We also classified simulated interactions into four epistasis types, based on the dose matrix response shapes (Fig 2C and D). These analyses were performed for all 1225 possible pairwise combinations of our 50 targets (S1 Table).

The single agent responses were very similar between methodologies, but combinations differed greatly (Fig 3 and S2-S4 Figs). Both FBA-res and FBA-div single agent activities were consistent with FBA knockouts (R~1, Fig 4). Moreover, even the IC50 concentrations showed a strong quantitative correlation (R=0.98, Fig 5). For combinations (Fig 4B), all three approaches mostly generated non-interacting pairs, with max(I) reflecting the more effective single agent’s, and no epistasis. However, there is a clear increase in the number and variety of interactions between FBA-ko, FBA-res, and FBA-div. While FBA-ko synergies were restricted to synthetic lethal (SL) epistasis, some partial synthetic lethal (PSL) appeared in FBA-res, and a large number of potentiation (Pot) synergies were added with FBA-div.

**Fig 3.**
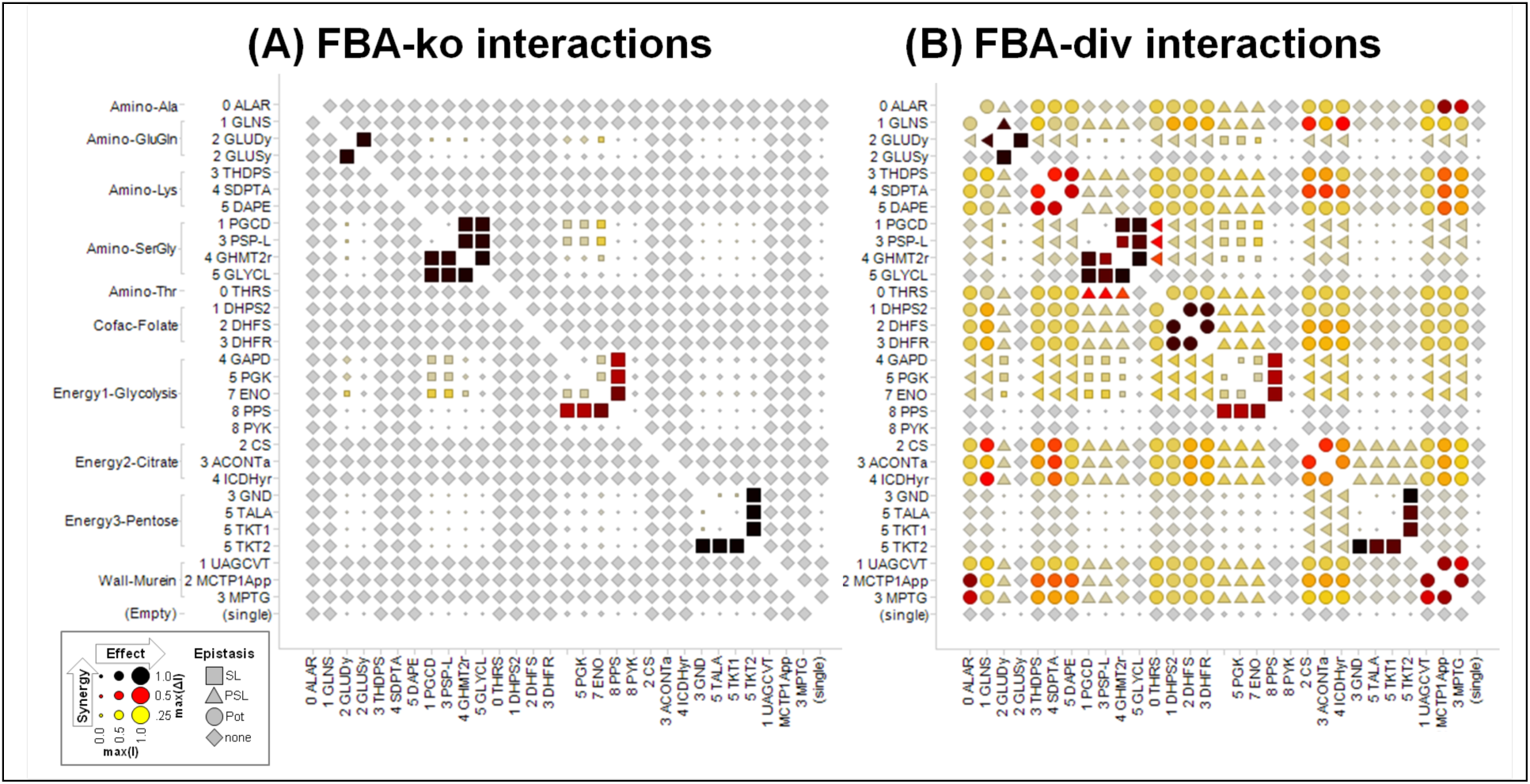
Comparing FBA-ko and FBA-div interactions across metabolic pathways. Simulated interactions for (A) FBA-ko and (B) for FBA-div, in selected metabolic pathways. More complete simulation results are shown elsewhere (S2-S4 Figs). Each symbol represents the simulated response to inhibiting a single or pair of targets (see Fig 2), with enzymes organized by pathway and metabolic reaction order in the iAF1260 model. Symbol shape shows the type of epistasis (synthetic lethal “SL”, partial synthetic lethal “PSL”, potentiation “Pot”, or non-interaction “None”), size shows the effect max(I), and color shows the synergy max(ΔI) between two targets (same scale as Fig 2). Single agents are shown along the bottom and right edges. The triangles for PSL epistasis point toward the inactive agent in the pairing. Single enzyme effects are very similar between FBA-ko and FBA-div, but interactions are very different. Most notably, strong Pot synergies are observed between serial targets in the same pathway under FBA-div (A), which are not predicted by FBA-ko (B) or FBA-res (S4 Fig).

**Fig 4.**
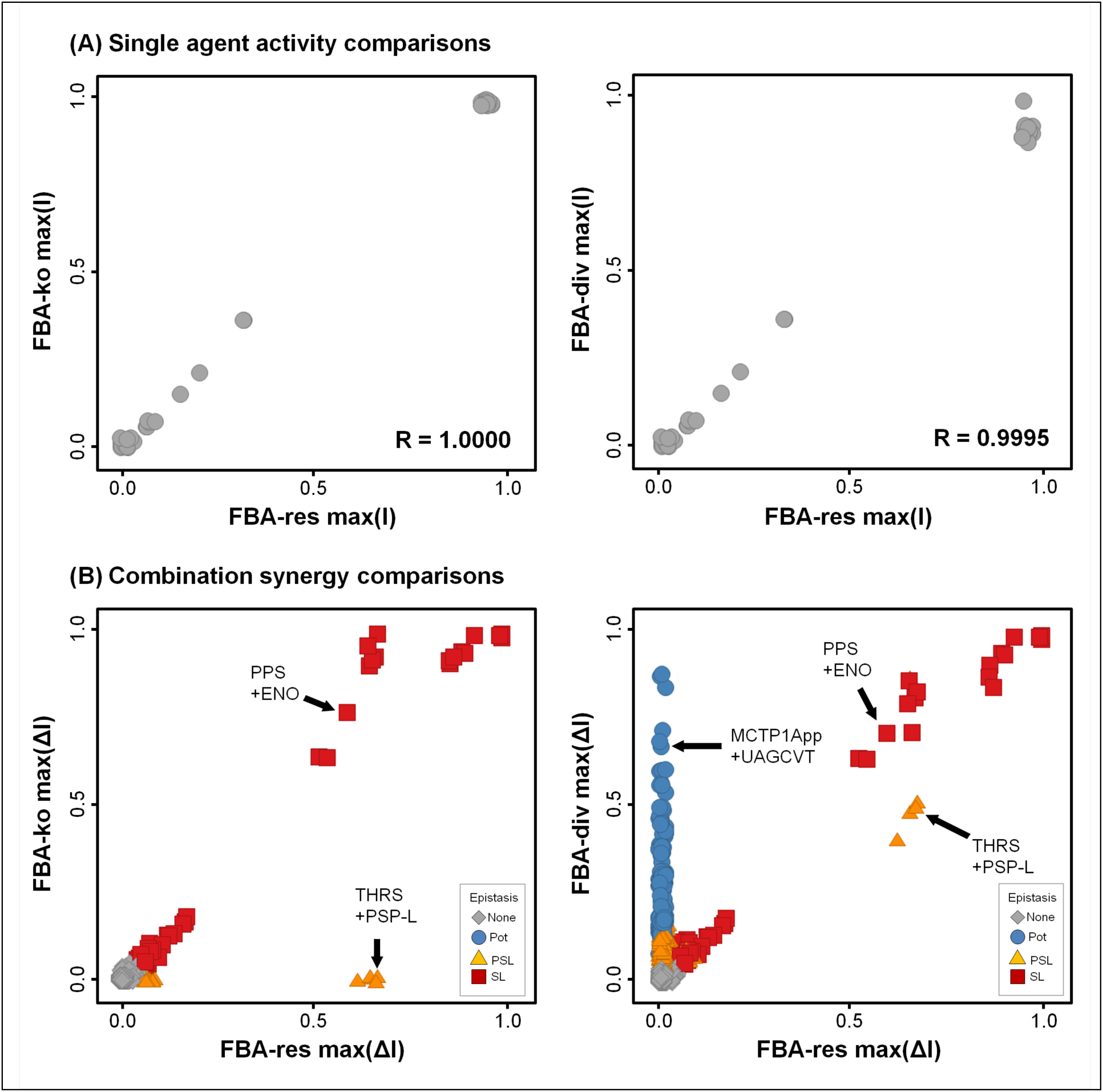
Comparing single agent and combination effects between methods. (**A**) Simulated single agent inhibitions are very consistent between methodologies. Across the 50 simulated inhibitors, both FBA-res and FBA-div methodologies yielded very similar max(I) scores to those obtained from standard FBA-ko. Data were randomly dithered by ~0.03 in both directions to visually separate overlapping points. (**B**) Simulated combination effects, however, show many deviations in max(ΔI) from FBA-ko, and substantial differences in epistasis types. Across all 1225 simulated pairs, FBA-ko yielded mostly None, with a few SL (2%). Simulating with FBA-res converted a few of the None to PSL (5%), and FBA-div reproduced all the FBA-res synergies and converted another 16% from None to Pot. Data were randomly dithered by ~0.03 to separate overlapping points.

**Fig 5.**
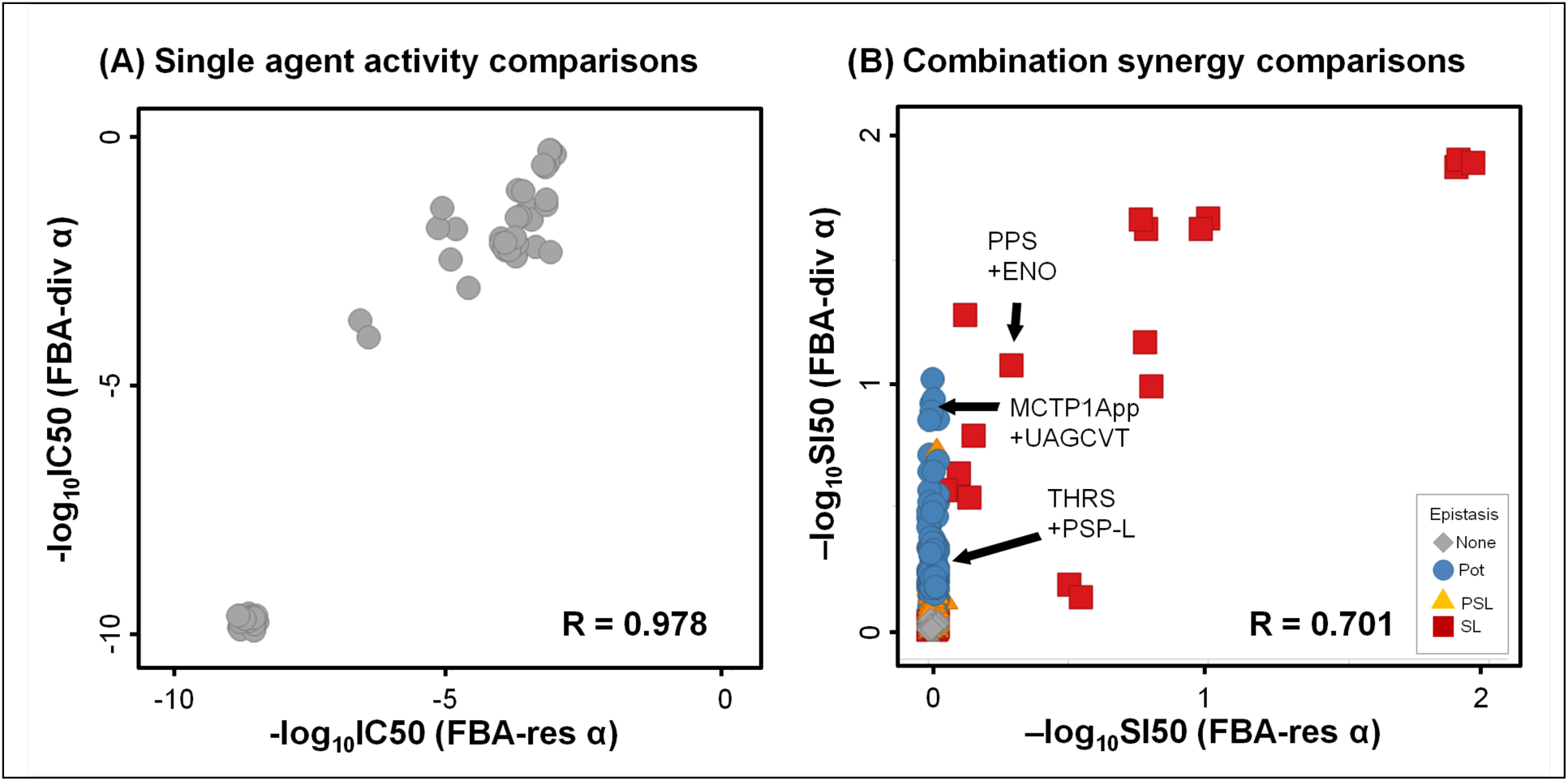
Comparing IC50 potency and SI50 synergy scores between simulation methods. (**A**) Simulated single agent inhibition potencies (as −logCI50) are very consistent between methodologies, though FBA-div responses are consistently more potent. Data were randomly dithered by ~0.3 in both directions to visually separate overlapping points. (**B**) Combination synergy, as measured by −logSI50, also is very consistent for non-interactions and synthetic lethalities, again with stronger synergies in FBA-div. Data were randomly dithered by ~0.03 in both directions to visually separate overlapping points.

Arranging the results by target pathway reveals informative patterns (Fig 3 and S2-S4 Figs). First, we located our iAF1260 targets on the iJR904 [37] pathway maps, and then assigned gene identities with reference to KEGG [34]. Pathways were then represented using concise descriptors listed in S2 Table. All three simulation approaches show active single agents clustered in pathways essential for *E. coli* growth in rich media (glycolysis, citrate cycle, folate and murein biosynthesis, and certain amino acid pathways), and the responses for any particular enzyme are more or less uniform within each such pathway.

Combination effects within pathways and interactions across pathways were mostly consistent across targets in each pathway even though they vary substantially between the different simulation methodologies. Specifically, FBA-ko (Fig 3 and S2 Fig) generated a small number of SL interactions, all of which are between individually inactive alternative reactions that are connected by a shared downstream output. Examples include TKT2 with other Energy/pentose enzymes, PPS with other Energy/glycolysis enzymes, and multiple isozyme pairs (GLUDy+GLUSy, TRPS1+TRPS3, GARFT+GART, DHORD2+DHORD5). There are also weaker interactions between PGL in Energy/pentose and the Energy/glycolysis enzymes, as well as between the glycolysis and Amino/SerGly pathways. Inhibiting with FBA-res (S3 Fig) reproduced all of these effects, but also added some PSL synergies involving one responsive target, either between closely connected enzyme pairs (eg, GLUDy+GLNS or between the Energy/pentose and Energy/citrate pathways). Finally, FBA-div simulations (Fig 3 and S4 Fig) recapitulated all of the FBA-ko and FBA-res effects, with a large number of additional Pot synergies. Generally, Pot occurred between pathways with active targets, especially between serial targets in the same pathway. Interestingly, the level of Pot synergy varied, with the strongest between serial targets in the same pathway (eg, in folate, murein, and amino-Lys). Cases where specific enzymes in a pathway stand out can be explained. For example in murein biosynthesis, there is strong FBA-res synergy between ALAR in amino-Ala with MCT1App and MPTG but the synergy is much weaker with UAGCVT (Fig 3 and S4 Fig). This can be understood as an extension of serial target synergy, because multiple alanine moieties dependent upon ALAR are added by MCT1App to its substrate, but below UAGCVT in the pathway. Another example is the especially strong interaction of SDPTA in amino/Lys with targets in Energy/citrate. A by-product of SDPTA is alpha-ketoglutarate (akg), a key metabolite in the citrate cycle.

### Diverted flux FBA predicts clinical drug synergies

To test how well each of the simulation methodologies models inhibited bacterial metabolism, we selected 28 combinations for experimental testing. Combinations were chosen to represent known synergistic and non-synergistic combinations, using compounds that sample different mechanisms of action across the metabolic network. The set includes cell wall metabolism inhibitors ampicillin, aztreonam and fosfomycin, folate synthesis inhibitors sulfamethoxazole and trimethoprim and metabolic inhibitors of central metabolic enzymes. The effect of each combination was tested in *E. coli* cultures using an 8×8 drug concentration-gradient matrix, with twofold dilutions between each step. Each top concentration was selected to capture published *in-vitro E. coli* responses, and combinations were tested at all possible pairs of each drug’s concentration series (S3 Table). Results are reported in S4 Table, with combination response matrices displayed in S5 Fig. Calculated response and synergy scores are integrated with the model scores (S1 Table).

Given the variety of pharmacodynamic effects that influence single agent potencies, we did not expect and also did not find strong agreement between the simulated and experimentally observed single agent max(I) or IC50 values (S1 Table). Aside from the clinical antibiotics, there are very few well characterized probes with selective activity on single metabolic targets, so it is not surprising that half of the compounds showed no activity against *E. coli* proliferation (S1 Table). The antibiotics showed some agreement with the FBA simulations in terms of Max(I) and IC50, but only in a qualitative sense.

For combinations, however, flux-diverted FBA was able to model the strong antibiotic synergies that target serial enzymes within a pathway (Fig 6A). FBA-div predicts the large potency shifts seen for both Sulfx+Trimp and Ampcl+Aztrm, while FBA-res and FBA-ko do not (Fig 6B and S1 Table). The experiments also found a number of moderate synergies (Fig 7), most of which involve inhibitors of targets just upstream of the same two pathways (eg, Ampcl or Aztrm combined with Fosfm or Cyser). Only a hint of the antibiotic synergy is detected for those probes, however, most likely due to less-specific enzyme targeting. Combination effects were absent, as expected, for those probes showing no single agent activity and thus not likely to have any relevant activity on the ascribed target. Finally, there was one experimental antagonism (Ampcl+Sulfx), that was not predicted by any of the FBA methods. Overall, comparing synergy scores (Fig 7), FBA-res and FBA-ko synergy scores shows no consistency with the experimental results, while the FBA-div simulations yield significant positive correlations (R ~ 0.44, p < 0.01) for max(ΔI). Accordingly, FBA-div represents the most accurate computational predictor of serial target anti-metabolic synergies.

**Fig 6.**
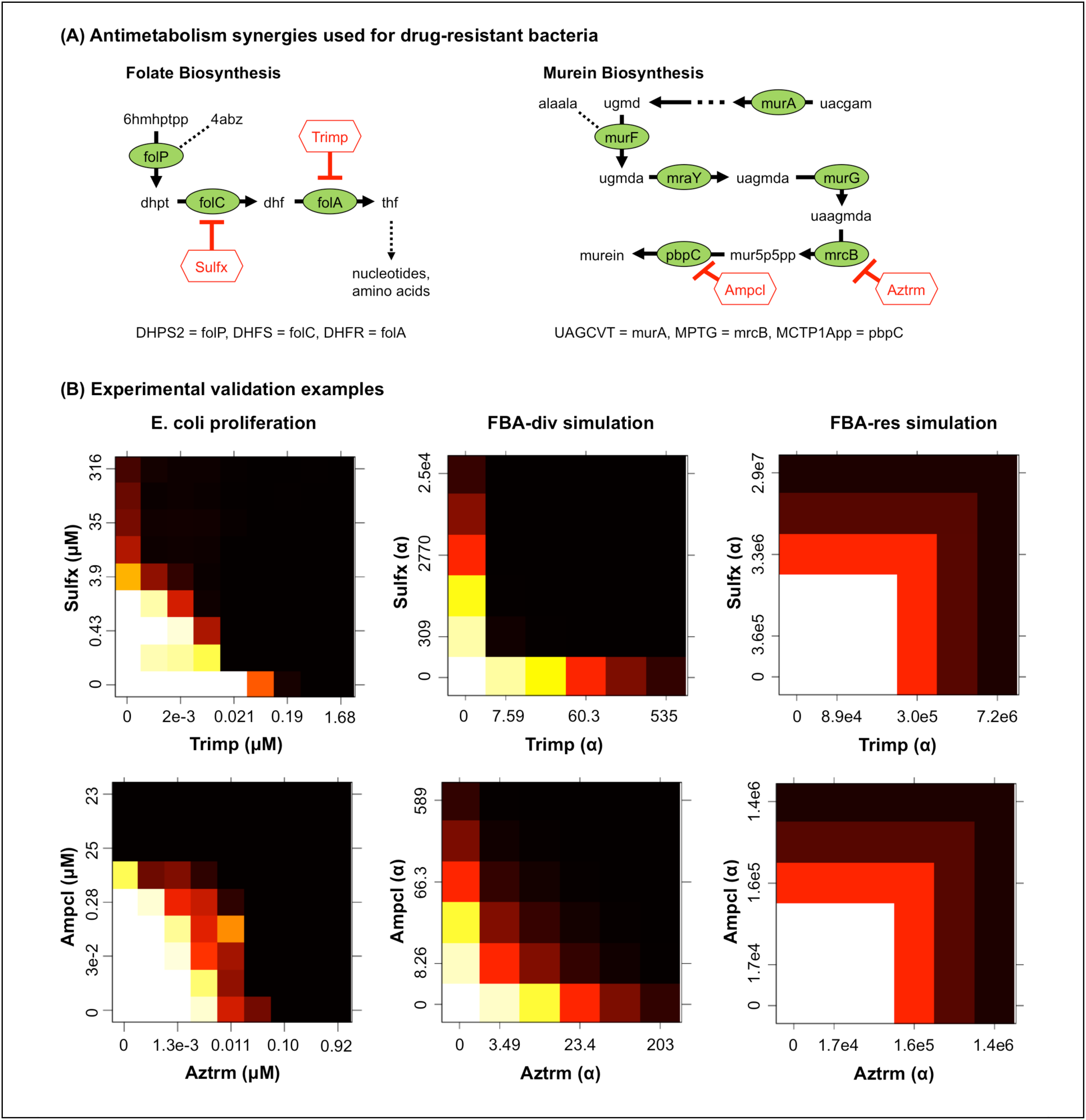
Experimental confirmation of simulated combination effects. (**A**) Known anti-metabolism drug synergies. Locating the iAF1260 model enzymes in the KEGG E. coli MG1655 pathway maps, sulfamethoxazole + trimethoprim inhibits folC+folA in folate biosynthesis, while aztreonam + ampicillin targets mrcB+pbpC in murein synthesis. We tested both combinations using an E. coli proliferation assay. (**B**) Response surfaces for the two known antibiotic combinations match the FBA-div simulations more closely than FBA-res. Although the models yield similar target inhibition levels (though requiring larger α for FBA-res), only FBA-div predicts the observed Pot synergy.

**Fig 7.**
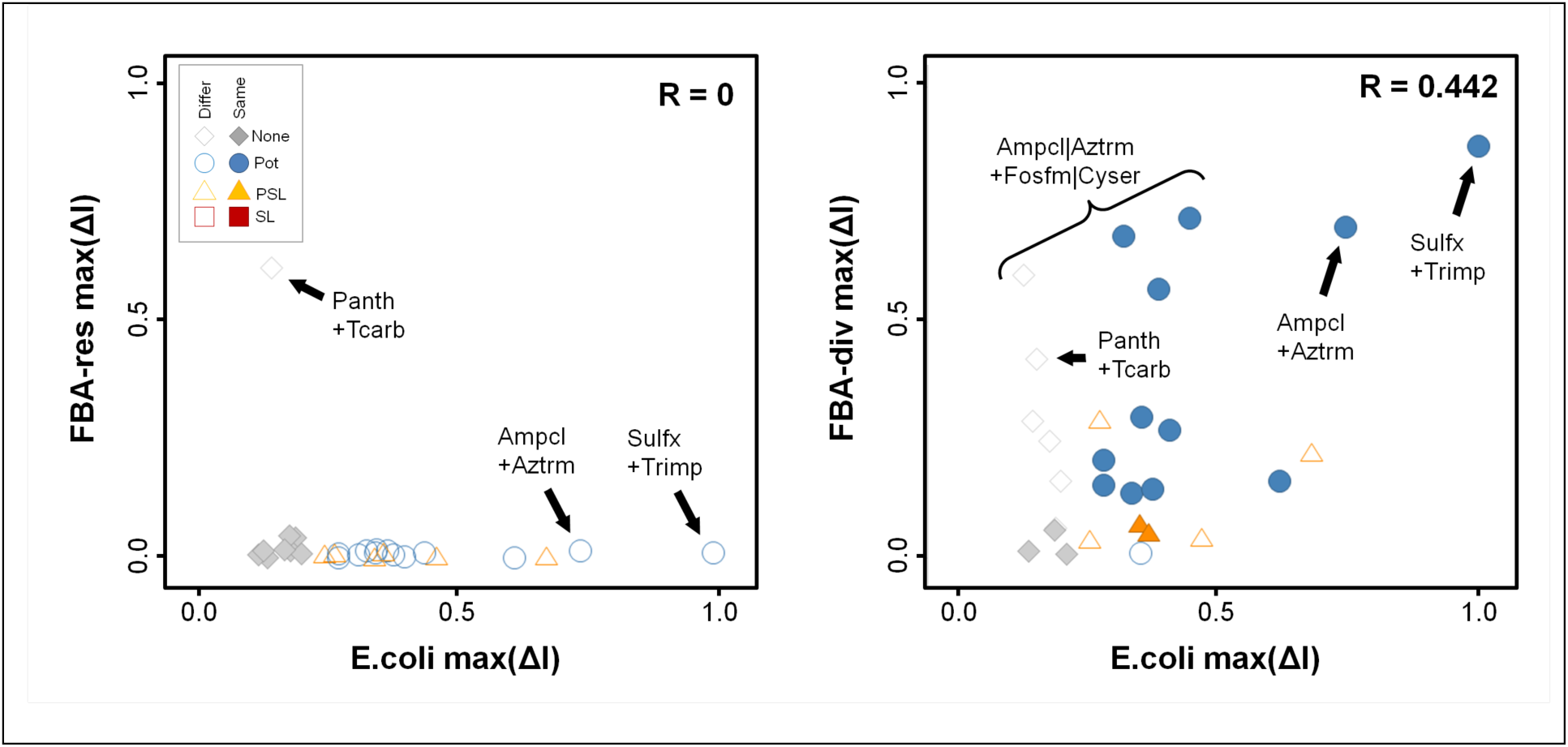
Experimental confirmation of simulated combination effects. We tested 28 combinations of metabolic inhibitors using an E. coli proliferation assay. Synergy score comparison for all combinations, where shape/color shows experimental interaction class, and open symbols indicate epistasis type. We find no agreement (R~0) between the experimentally determined drug interactions and FBA-res. However, experimental and FBA-div synergy are correlated (R~0.44). In addition to the antibiotic combinations, weaker synergy is predicted and observed for other interactions between murein synthesis inhibitors and targets further upstream.

## Discussion

While FBA methodologies have shown great potential for rationalizing metabolic engineering efforts and identifying synthetic lethal genetic dependencies, traditional FBA knockout simulations fail to predict the most useful antibiotic synergies that target metabolism. As drug combination therapies gain importance for antibiotic treatment, more accurate large-scale prediction methods are sorely in demand. Here we show that extending FBA simulation using flux diversion can accurately model the strong dose-dependent synergies observed between inhibitors of bacterial metabolism.

The FBA-div methodology outperforms both FBA-res and standard knockout FBA for predicting experimental combination effects, especially for one of the most important antibacterial combinations that target metabolism. Prior approaches for simulating drug epistasis completely missed synergies like sulfamethoxazole and trimethoprim, because knockout and restriction-based approaches find no interactions between serial targets in a pathway. In contrast, FBA-div predicts strong synergies from targeting sequential targets in the same pathway, similar to what has been found for competitive inhibition of serial targets with negative feedback in kinetic simulations[10]. While this is not the usual expectation from paired knockout analyses[18] it is consistent with strong antibiotic synergies being used in the clinic. Among our 28 tested combinations, the correlation between FBA-div and experimental synergy scores is modest (R ~ 0.442), due to the scarcity of selective metabolic inhibitors. However, with metabolism increasingly a focus of drug development, especially for combinations, FBA-div simulations could help discover and prioritize drug targets in multiple disease areas.

Broader patterns of synergy across pathways (eg, Fig 3 and S4 Fig) can also provide key insights into the functional connections that are most relevant to the system under study[10,18]. It is notable that there are consistent patterns of single agent activity and combination epistasis within a pathway, and again consistent patterns between targets across pathways, confirming target connection topology as a major determinant of simulated combination effects[10,18]. All three methods discussed here find strong synthetic lethality between isozymes of essential reactions, and weaker interactions between alternative essential pathways. However, FBA-div provides by far the richest source of predicted synergies. For example, the citrate cycle synergizes with Lysine metabolism, but exceptionally strongly with SDPTA, revealing a direct connection through that enzyme’s by-product alpha-ketoglutarate. Similarly, other connections with amino acid metabolism are revealed for the murein and folate biosynthesis pathways.

Differences between the FBA-res and FBA-div methodologies may be understood in terms of standard Michaelis-Menten (MM) modeling for drug-inhibited enzymes. In MM kinetics, the reaction velocity **v** = V_M_**S**/(**S**+K_M_), where **S** is the input substrate’s concentration, K_M_ is the reaction affinity, and V_M_ is the maximum reaction rate. In FBA-res, a scalar reduction in **v** by α corresponds to a perturbed 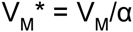, while 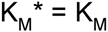, since the stoichiometric parameters **s**_ij_ that account for reaction affinities are unchanged. This is the behavior of non-competitive inhibitors in MM kinetics with α=(1+[drug]/K_D_). By contrast, in FBA-div, scaling down the reaction affinity corresponds to 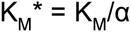, this time leaving 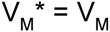 constant, similar to competitive inhibitors. These scaling behaviors are consistent with combination effects seen in kinetic pathway simulations[10], where competitive inhibition of serial targets with negative feedback yielded strong serial-target synergies that were absent in simulations of non-competitive or uncompetitive MM reactions. Thus, FBA-div more closely mimics the inhibitory kinetics of metabolic inhibitors used as antibiotic drugs.

From a biology perspective, FBA-div can explain serial target synergies through the build-up of metabolite concentrations. The waste stream in FBA-div can be thought of as representing passive regulation that limits substrate concentrations by degradation or export from the system. Simulating with FBA-res assumes that when a drug inhibits a reaction, accumulated mass will slow down the upstream reaction thermodynamically, or induce a gene expression cascade that causes upstream reactions to slow down to the inhibited level. By contrast, FBA-div assumes that upstream reactions cannot sense or adjust to an inhibited downstream reaction on the time scale of the drug inhibition, and substrate is wasted, creating less biomass over time. Also, diverting excess flux to waste mimics processes similar to the phenomenon of metabolic resistance[25,38] which has been experimentally shown to result from upstream substrate accumulation. Such mechanisms have been proposed to explain how microbes keep substrate concentrations within feasible ranges, and FBA models have indeed been augmented with explicit constraints on metabolite and Gibbs free energy levels to more accurately predict global flux distributions[39]. Put in simpler terms, FBA-res cannot predict serial target synergies because mass conservation during optimization requires any flux restriction to apply throughout a serial pathway, forcing the combined effect to match that of the most effective inhibitor. In FBA-div, however, inhibition diverts intermediate reaction flux to waste, yielding greater biomass reductions.

The FBA-div approach can be extended to metabolic models of pathogens, such as tuberculosis[40], plasmodium[41], and the ESKAPE pathogens[42]. As more FBA models become available for pathogens, combination effects within the current space of drugs and drug-like small molecules could be simulated, and new therapies could be rapidly tested against drug-resistant strains. Indeed, FBA-div for human metabolism is increasingly conceivable as mammalian FBA models[43] become more established. Our FBA-div perturbation approach could also be used with simulation methodologies that extend FBA to transcriptional regulation using ChIP-Seq and gene expression information[44], enabling yet more realistic simulations of drug effects in biological systems. Finally, flux diversion may also more accurately predict synergies between 3 or more drugs[4,13,45]. These extensions would also enable useful synergy predictions for drug combinations targeting mammalian metabolism.

Experimental work on drug combinations can be resource intensive, especially for pathogens which require a BSL 3 environment, or when large combinatorial spaces are comprehensively sampled. Thus, computational tools that can explore large-scale models to prioritize such resource-intensive work could significantly accelerate drug discovery research. Because it can accurately predict a great variety of drug synergies, FBA-div has the potential to make a major impact on these therapeutic challenges.

